# Arm movements increase acoustic markers of expiratory flow

**DOI:** 10.1101/2024.01.08.574611

**Authors:** Raphael Werner, Luc Selen, Wim Pouw

## Abstract

The gesture-speech physics theory suggests that there are biomechanical interactions of the voice with the whole body, driving speech to align fluctuations in loudness and F0 with upper-limb movement. This exploratory study offers a possible falsification of the gesture-speech physics theory, which would predict effects of upper-limb movement on voice as well as respiration. We therefore investigate co-movement expiration. Seventeen participants were asked to produce a continuous exhalation for several seconds. After 3s, they execute one of five within-subject movement conditions with their arm with and without a wrist weight (no movement, elbow flexion, elbow extension, internal arm rotation, external arm rotation). We analyzed the smoothed amplitude envelope of the acoustic signal in relation to arm movement. Compared to no movement, all four movements lead to higher positive peaks in the amplitude peaks, while weight did not influence the amplitude. We also found that across movement conditions, positive amplitude peaks are structurally timed relative to peaks in kinematics (speed, acceleration). We conclude that the reason why upper-limb movements affect voice loudness is still best understood through gesture-speech physics theory, where upper-limb movements affect the voice directly by modulating sub-glottal pressures. Multimodal prosody is therefore partly literally embodied.

## 1. Introduction

Hand gestures are frequent during speech production. Often, these upper-limb movements have a pulsing ‘beat-like’ quality, characterized by sudden stops and accelerations temporally aligned with changes in voice intensity and fundamental frequency, two important acoustic correlates of stress [1]. The link between speech and gesture has been the subject of multi-modal prosody research [2]. Previous studies have found an effect of upper-limb movements on the acoustics for continuous voicing [3, 4], mono-syllable utterances [5], singing [6], and fluent speech production [7]. However, only movements of body segments with relatively higher mass and a beat-like quality incur effects on the voice, as continuous non-restive arm cycling does not seem to affect the voice [8], and beat-like movements of the wrist incur less effects on the voice than higher motion of proximal segments [3, 4]. Here we tackle another important issue in this line of research, raised by [9]: If the voice is affected by gesture through interaction with the respiratory system, as has been implied by research involving direct measurements of rib-cage movement [5] and as suggested by the gesture-speech physics theory [3, 10], then we should expect that expiratory rate would increase during upper-limb movement, even without vocalization. If not, the gesture-speech physics theory is wrong, and the effects it tries to explain should actually be understood solely through laryngeal and supra-laryngeal control.

Human speech production relies on respiration [11]. Breathing is not just affected by the *primary* respiratory muscles, such as the diaphragm or the intercostals [12], but also by *secondary* muscles that either activate during extreme respiratory conditions or during gesturing [13, 14, 15]. Furthermore any muscle that can affect the movement of the rib-cage can affect sub-glottal pressures and voicing, which is a larger set of muscles than those classically listed as primary/secondary respiratory muscles [12]. A recent study, directly looking at muscle activations, was the first to show that gesture kinematics of the upper body materialize as amplitude peaks in sustained vocalizations [16]. The analyses further showed that the voice interacted with whole-body muscle activity (measured via surface EMG and postural measurements), as well as mass of the movement (via wrist weights). Thereby weighted movements increased coupling of gestures and voice as compare to non-weighted movement.

So how can upper-limb movement affect the rib cage? According to gesture-speech physics theory there are two routes [10]: The first route is when a muscle is activated that moves the arm or stabilizes the arm movement, which also happens to be a muscle that inserts or indirectly affects rib cage movement. There are many such arm movements: for example an internal rotation of the humerus requires the activation of the pectoralis major (a chest muscle), which directly attaches to the rib cage. To move the arm, then means, to affect in some way rib cage movement too (according to the gesture-speech physics theory). Direct evidence for this first route has been found by pectoralis major activity predicting voice peaks [16].

The second route is through anticipatory postural adjustments that react to postural perturbations that are caused by beat-like rapid upper-limb movements. For example it has been shown that when the upper limbs are moved quickly, secondary respiratory muscles such as the abs activate [17], and even the diaphragm has been shown to stabilize the trunk and tension as a way to increase postural stability during upper-limb movement [13, 18]. Thus, when moving the upper limbs, just before and after (about *±* 50ms [14]) a peak acceleration or deceleration of the arm, one might expect other posture-stabilizing muscles to activate that directly affect rib-cage movement, too. Indirect evidence of this second route has been found in research showing that gesture-vocal coupling is more pronounced in a standing versus sitting position [3], and direct evidence was obtained that postural muscle in the back called the erector spinae was activated during upper-limb movement and also predicted amplitude peaks in the voice [16].

Importantly, despite the gesture-speech physics theory clearly assuming a respiratory effect of beat-like upper-limb movements, no measurements have been taken to directly assess respiratory modulation. In this exploratory study, we thus analyze the data of participants producing voiceless expirations that were recorded along with and under the same experimental conditions as voicing [16, 19]. We assess whether different types of arm movement affect expiration as compared to a passive no movement condition. We also assess whether particular movements increase expiration more than other types of movements. We will not measure expiratory rate directly, but the acoustic envelope. When air flow increases and moves against an object such as the lips and the microphone, a sound is produced with a certain intensity proportional to the rate of flow.

We mainly explore two questions here: 1) Do particular movement conditions predict stronger negative and positive peaks in amplitude envelope? 2) Are the positive/negative peaks observed in the amplitude envelope structurally related to the movement?

Following the findings from [16], we assume for 1) that similar to voicing, we should find increased positive peaks in the amplitude envelope reflecting expiratory rate. Negative peaks may be observed, but with less clear structural relations with movement. Further, since the acoustic intensity of expiratory sounds is much smaller in magnitude as compared to voicing, we anticipated that the effects of upper-limb movement will be harder to detect. If we measured expiratory rate more directly through spirometry, however, we would not predict such diminished effects. So if we find an effect in acoustics of expiration we believe we can make a good case that significant effects on respiration are to be expected (which warrants a further confirmatory study using spirometry). Regarding 2), [16] showed that only positive peaks were consistently found at particular moments of the movement (close to the acceleration phase of the movement). Here we expect a similar distribution for positive and negative peaks, namely that positive peaks will be more affected by upper-limb movement and thus show higher probability density at particular moments of the movement. We predict more random observations of negative or positive peaks in acoustics during the no movement condition.

## 2. Methods

The experimental setup is analogous to [16], where we analyzed sustained vocalizations. 17 participants (7f, 10m) were asked to produce continuous exhalation or vocalization with their elbow flexed at a 90° angle^1^. We here only include the exhalation condition. After 3 s, they were prompted to execute one of five movement conditions with their arm (no movement, elbow flexion, elbow extension, shoulder internal rotation, shoulder external rotation), while keeping the exhalation as steady as possible for another 4 s. The movement conditions are depicted in Fig. 1. For the no movement condition participants were asked to rest their arms alongside their bodies. Additionally, there was a within-subject weight condition to manipulate the mass set in motion, i.e. participants either wore a 1 kg wrist weight or nothing at their wrist.

**Figure 1.**
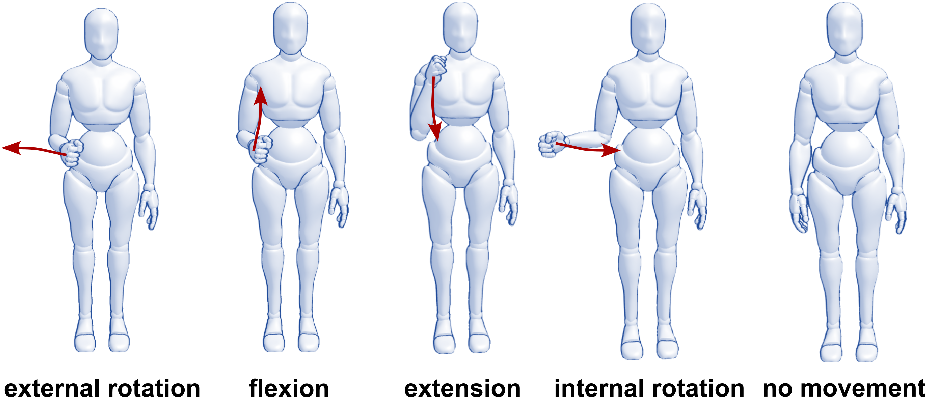
The five arm movement conditions.

The experiment was coded in Python using functions from PsychoPy. We recorded audio (via a head-mounted microphone with fixed gain levels), video for 3D motion tracking, muscle activity (via surface electromyography) of four muscles involved in rotation of the arm or posture (pectoralis major, infraspinatus, erector spinae, rectus abdominis), as well as postural perturbation via a balance board. We exclude posture and muscle activity from the analyses here. Still, the activity of those muscles is indirectly included, as there is a correspondence between the movements and a particular muscle. That is, pectoralis major drives internal rotation, while the infraspinatus drives external rotation. Rectus abdominis and erector spinae are postural muscles.

For data synchronization, we used LabStreamingLayer^2^. All signals were sampled at, or upsampled to, 2,500 Hz. For the video data, we used Mediapipe [20] to track the skeleton and facial movements, which is implemented in Masked-piper which we also use for masking the videos [21]. The video-based motion tracking is used to determine the the peak speed of movement, and then locate other signal peaks around them (and to centralize timeseries relative to the kinematic landmarks, such as peak speed, or movement onset). For the audio signal, we extracted the smoothed amplitude envelope. For acoustics, we measured the global maxima happening within the analysis window, i.e. between movement onset and offset. We also analyze global minima separately. To cater for de- or increasing amplitude over the exhalation, we de-trend this line and assess positive/negative peaks relative to this trend line. Note, that the participants are asked to keep expiratory flow as steady as possible, so we measure deviations from steady flow with this detrending approach. As the distribution of positive and amplitude peaks had long tails, we log-transformed them before running the statistical models. In the case of negative peaks, we absolutized the values before the transformation (as you cannot take a logarithm of negative values). Note that we excluded trials where we did not obtain a negative or positive peak relative to trendline (we excluded N = 17 data points from positive peaks, i.e. 2.7%, and we excluded N = 32 from negative peaks, i.e. 5.1%).

## 3. Results

We first focus on question 1), i.e. do particular movement conditions result in stronger negative and positive peaks in the amplitude envelope? The smoothed amplitude envelopes grouped by movement condition and centered at movement onset can be seen in Fig. 2. From the envelope time series underlying these smoothed trajectories we extract positive and negative peaks in a window from 500 ms before to 500 ms after the movement onset/offset (using peakfinding function on the 2D speed time series of the wrist). By positive peaks we refer to the highest points on the vertical axis, by negative peaks to the lowest and typically negative values. This visualization already suggests that the movement conditions have different effects on the exhalation amplitude than the no movement condition. All movements lead to clear positive peaks, while negative peaks are not as clear in all movement conditions.

**Figure 2.**
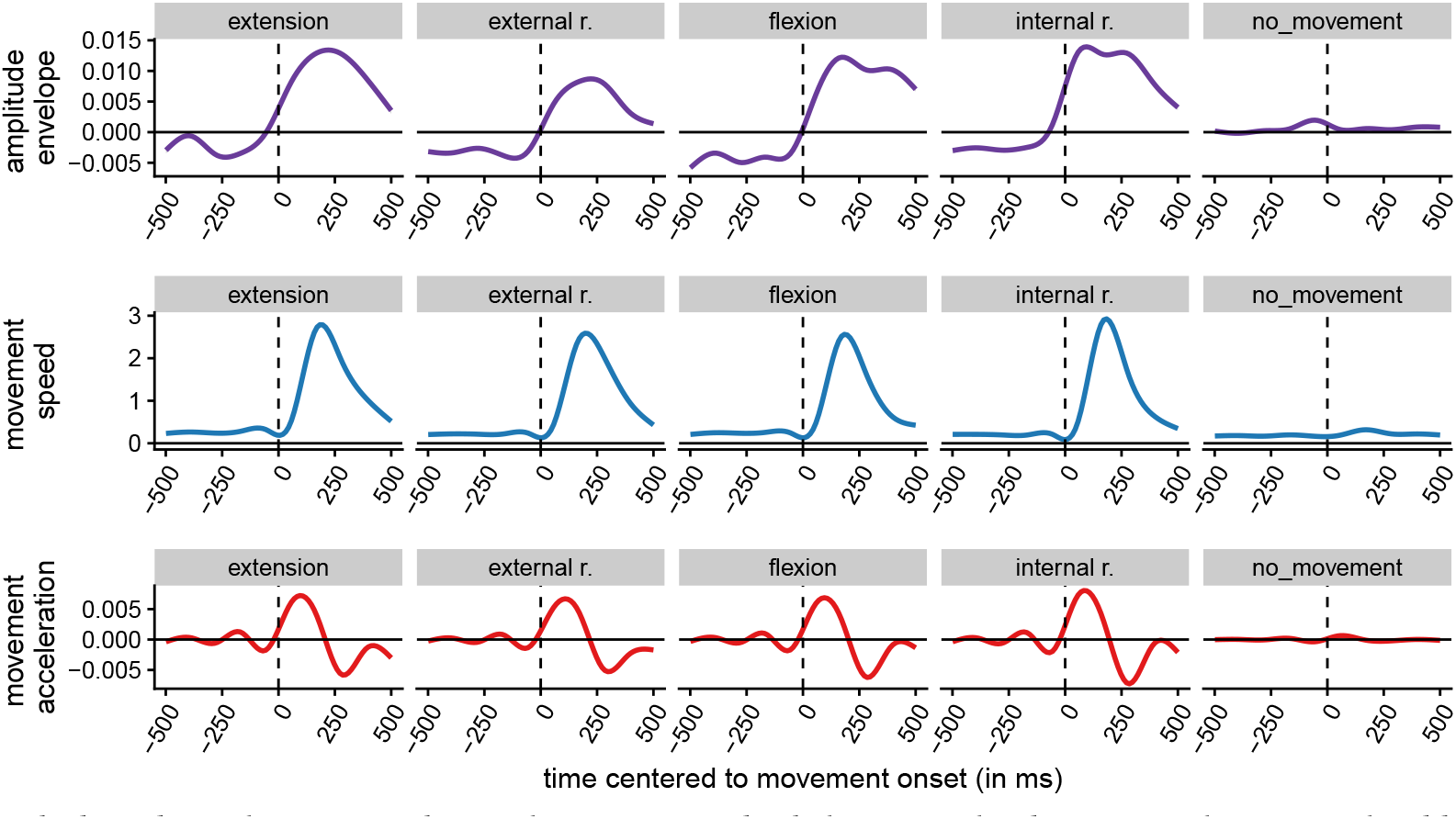
Smoothed conditional means for the non-log-transformed exhalation amplitude (top, purple; z-normalized by participant), movement speed (middle, blue; z-normalized by participant), and movement acceleration (bottom, red; derivative of z-normalized speed) by movement condition across participants (via fitting non-linear smooths using generalized additive models in R-package `ggplot2`). Time zero is defined as movement onset (as determined by the motion tracking of the wrist).

The log-transformed amplitude peaks, both positive and negative, can be seen in Fig. 3, split by movement and weight conditions. Using R-package `lme4`, we modeled the variation in positive amplitude peaks by weight and movement condition, with participant as random intercept. A model with the movement and weight conditions explained more variance than a base model predicting the overall mean, change in Chi-Squared (5.00) = 19.01, p = 0.002. Model results can be found in Table 1. Compared to exhaling with no movement, all movement conditions had a positive effect on positive amplitude peaks. Adding weight to the wrist did not influence the amplitude in this model. As the no movement condition is not affected by weight, we ran another model, where we excluded the no movement control condition. This model, with only weight as a fixed effect and participant as random intercept, also, shows that weight did not have a significant effect on the voice’s positive peaks (b = 0.032, SE = 0.073, t = 0.439, p = 0.661).

**Figure 3.**
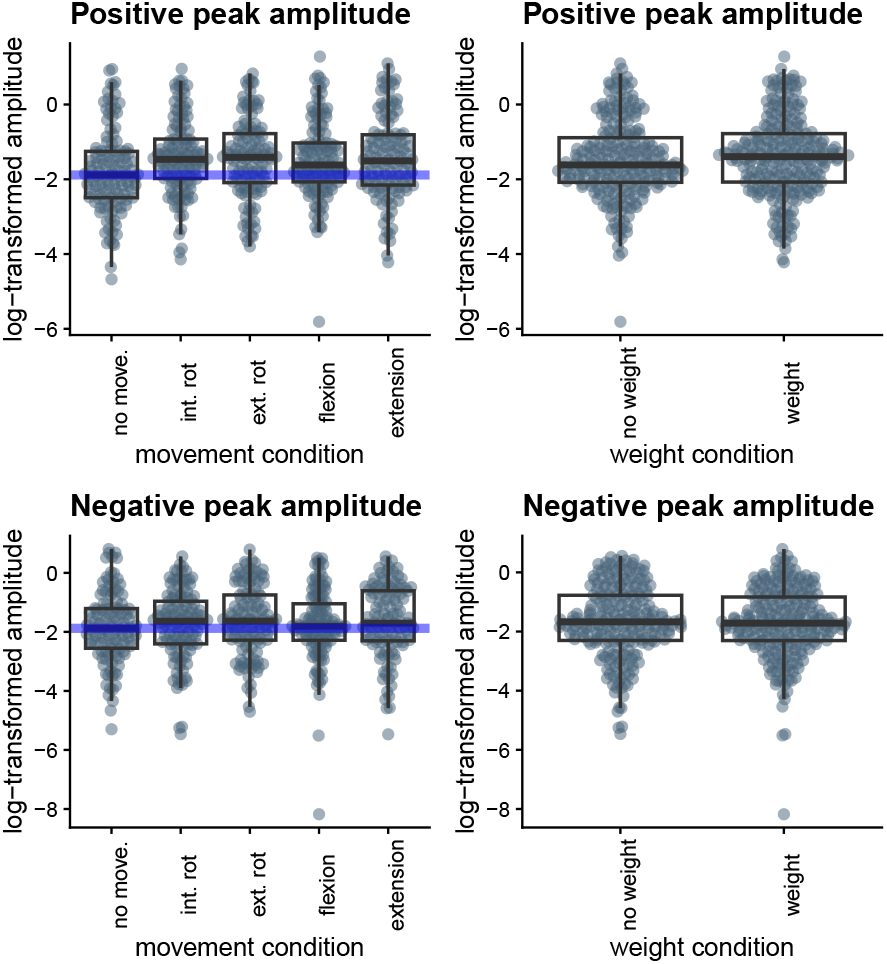
Box plots showing log-transformed, positive (top) and negative (bottom) peaks in the amplitude envelope. The higher the values, the higher the magnitude of the positive or negative deviation. The panels on the left show the effects by movement condition, the panels on the right show them by weight condition. Blue lines show the median of the no movement condition in the respective panel.

**Table 1:** Results of the model predicting positive amplitude peaks by weight and movement conditions.

|  | Estimate | SE | df | t-value | p-value |
| --- | --- | --- | --- | --- | --- |
| Intercept (no movement/no weight) | -1.781 | 0.190 | 22.331 | -9.356 | < .001 |
| vs. weight | 0.062 | 0.066 | 586.558 | 0.933 | 0.351 |
| vs. internal rotation | 0.395 | 0.104 | 586.024 | 3.786 | < .001 |
| vs. external rotation | 0.344 | 0.105 | 586.045 | 3.268 | 0.001 |
| vs. flexion | 0.303 | 0.104 | 586.024 | 2.905 | 0.004 |
| vs. extension | 0.342 | 0.105 | 586.048 | 3.269 | 0.001 |

We then modeled the negative peaks by weight and movement conditions, with participant as random intercept. We found no significant effects on negative amplitude peaks, neither for weight or any of the movement conditions. Neither a model with movement conditions, weight and movement conditions, or interactions of both was able to explain more variance than a base model predicting the overall mean. For the additive model, we found change in Chi-Squared (5.00) = 2.94, p = 0.709. The absence of weight condition effect did not change when including only movement conditions. This means that the negative peaks of the exhalation amplitude were not affected by any of the movement or weight conditions, when compared to the no movement and no weight condition.

Finally, we addressed question 2) about timing between positive and negative peaks in the amplitude envelope relative to peak speed of hand movement. Fig. 4 shows the temporal relationships of negative and positive peaks in relation to when the hand reaches peak speed at t = 0 (please note: we change alignment here from onset to peak speed). In the timing relations, the negative amplitude peaks are more evenly distributed in the time window peak speed, while for the positive peaks there are clear occurrences just before the peak in speed. Note we only look at peaks before peak in speed (otherwise peaks would also show up at deceleration). The no movement conditions (solid lines) further suggest that for the expiration in this experiment, different types of upper-limb movement did have an effect on the occurrence of positive, but not for negative amplitude peaks.

**Figure 4.**
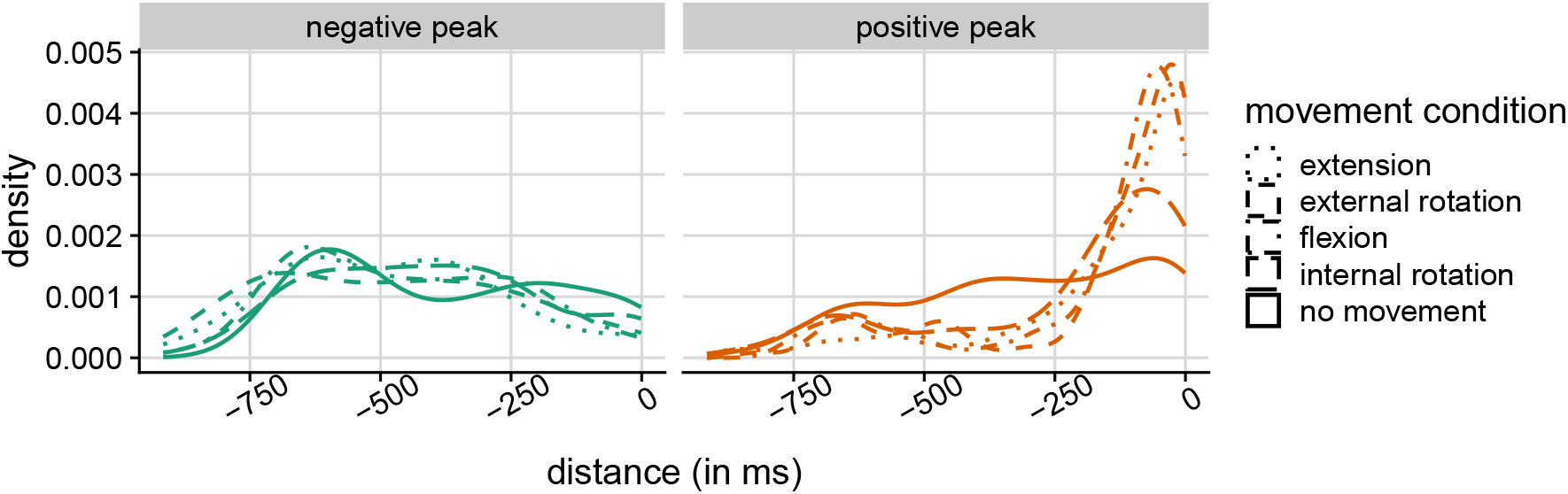
Density plot showing temporal distance (in milliseconds) of negative (left) and positive (right) peaks in amplitude relative to peak speed in wrist movement (at t = 0).

## 4. Discussion

In this study we find effects of different movements on positive amplitude peaks in the acoustics as a measure of flow of voiceless expiration. We did not find an effect for negative peaks, or an impact of adding weight to the wrist in any condition here. In the vocalizations [16], movement and weight did affect both, positive and negative peaks of the vocal amplitude, though only positive peaks seemed to be structurally driven by particular moments in the movement. For expiration we also find that the temporal relation between positive peaks in expiration amplitude and hand movement were very similar to those found for vocalizations [16]. The current results suggest that the effect of upper-limb movement on voicing is indeed driven by the movement affecting subglottal pressure and thus expiration, and cannot be understood only through (supra-)laryngeal modulation. It is further important to underline that since acoustic measurements of expiratory noises are not direct measurements of expiratory flow, it is not surprising that we obtain lower effect sizes of movement on expirations (and no replication of the weight condition) as compared to voicing.

By disentangling respiratory modulation from voicing, as suggested by [9], we obtain a more direct test of the gesture-speech physics theory. We conclude that, so far, and even with imperfect measurement of respiratory modulation through acoustics, gesture-speech physics theory seems to hold [22]. Upper-limb movement during voicing or expiration produces modulations at particular moments of the movement that the theory predicts to be related to direct and indirect muscle actions that move the arm, or stabilize posture, during arm movement [10].

## 5. Conclusions

This exploratory study has shed light on the influence of gestures on vocal production via exhalation by analyzing positive and negative peaks in the exhalation amplitude envelope in relation to upper-limb movements. By taking voicing out of the equation, we corroborate the findings of [16] and show that these effects are driven by changes in subglottal pressure, rather than laryngeal modifications. However, the effects are less strong and clear in exhalations compared to vocalizations. While we were able to predict positive peaks in the exhalation amplitude envelope from the movement conditions, the picture was less clear for muscles and we found no effects for negative peaks anywhere. We thus conclude that upper-limb movement affects not just voiced speech production, but also exhalation.

This work adds to our understanding of multi-modal prosody, by strengthening the mechanistic understanding of the link of production between gesture and speech. In general the current work contributes to a wider investigation of how sounds produced by the body can be a source of bodily and correlated psychological states and processes [23, 24]. It also contributes to an understanding of language production as inherently entangled with *literal* bodily processes [25, 26, 27]. Not bodily processes that are embrained as some forms of embodied cognition have it [28], but literal physical processes that can become part of particular modes of speaking as more ‘radically embodied’ views on cognition and language would have it [25, 26, 27] (see also [29]). Moreover, the current research shows that breath noises not only inform about the speaker’s vocal tract configuration [30], but also their gestures - there is thus much more signal in ‘breathing noises’ than is typically assumed.

## 6. Acknowledgements

We would like to thank Pascal de Water at the Donders Institute for the formidable technical support.

## Footnotes

1 More details about the experiment can be found via <link redacted for review>.

2 See https://github.com/sccn/labstreaminglayer.

